# Kinematics of the Lever Arm Swing in Myosin VI

**DOI:** 10.1101/078626

**Authors:** M. L. Mugnai, D. Thirumalai

**Affiliations:** Biophysics Program, Institute for Physical Science and Technology, University of Maryland, College Park, MD 20742; Department of Chemistry and Biochemistry, University of Maryland, College Park, MD 20742

## Abstract

Myosin VI (MVI) is the only known member of the myosin superfamily that, upon dimerization, walks processively towards the pointed end of the actin filament. The leading head of the dimer directs the trailing head forward with a power stroke, a conformational change of the motor domain exaggerated by the lever arm. Using a new coarse-grained model for the power stroke of a single MVI, we provide the molecular basis for its motility. We show that the power stroke occurs in two major steps: first, the motor domain attains the post-stroke conformation without directing the lever arm forward; second, the lever arm reaches the post-stroke orientation by undergoing a rotational diffusion. From the analysis of the trajectories, we discover that the potential that directs the rotating lever arm towards the post-stroke conformation is almost flat, implying that the lever arm rotation is mostly un-coupled from the motor domain. Because a backward load comparable with the largest inter-head tension in a MVI dimer prevents the rotation of the lever arm, our model suggests that the leading-head lever arm of a MVI dimer is uncoupled, in accord with the inference drawn from polarized Total Internal Reflection Fluorescence (polTIRF) experiments. Our simulations are in quantitative agreement with polTIRF experiments, which validates our structural insights. Finally, we discuss the implications of our model in explaining the broad step-size distribution of MVI stepping pattern, and we make testable predictions.

## Introduction

Like their counterparts dyneins and kinesins, myosins are molecular motors that transform the chemical energy harvested in the hydrolysis of ATP into mechanical work. They do so by undergoing a reaction cycle (Fig. 1) involving ATP hydrolysis coupled with binding to and unbinding from the filamentous actin (F-actin) [1, 2]. Myosins loaded with a newly hydrolyzed ATP bind to F-actin, release the products of ATP hydrolysis, and undergo a structural change known as power stroke. The structural change of the N-terminal part of the motor domain, where the actin and nucleotide binding sites reside, is communicated to the converter domain that moves from the pre-power stroke (PrePS) state to post-power stroke (or Rigor, R) state conformation. The movement of the converter is exaggerated by the large swing of the lever arm, an oblong domain bound to light chains or calmodulins (CaMs). When a new ATP molecule binds the nucleotide-free myosin (in R state), the motor detaches from actin and it is ready to begin a new cycle.

**Figure 1:**
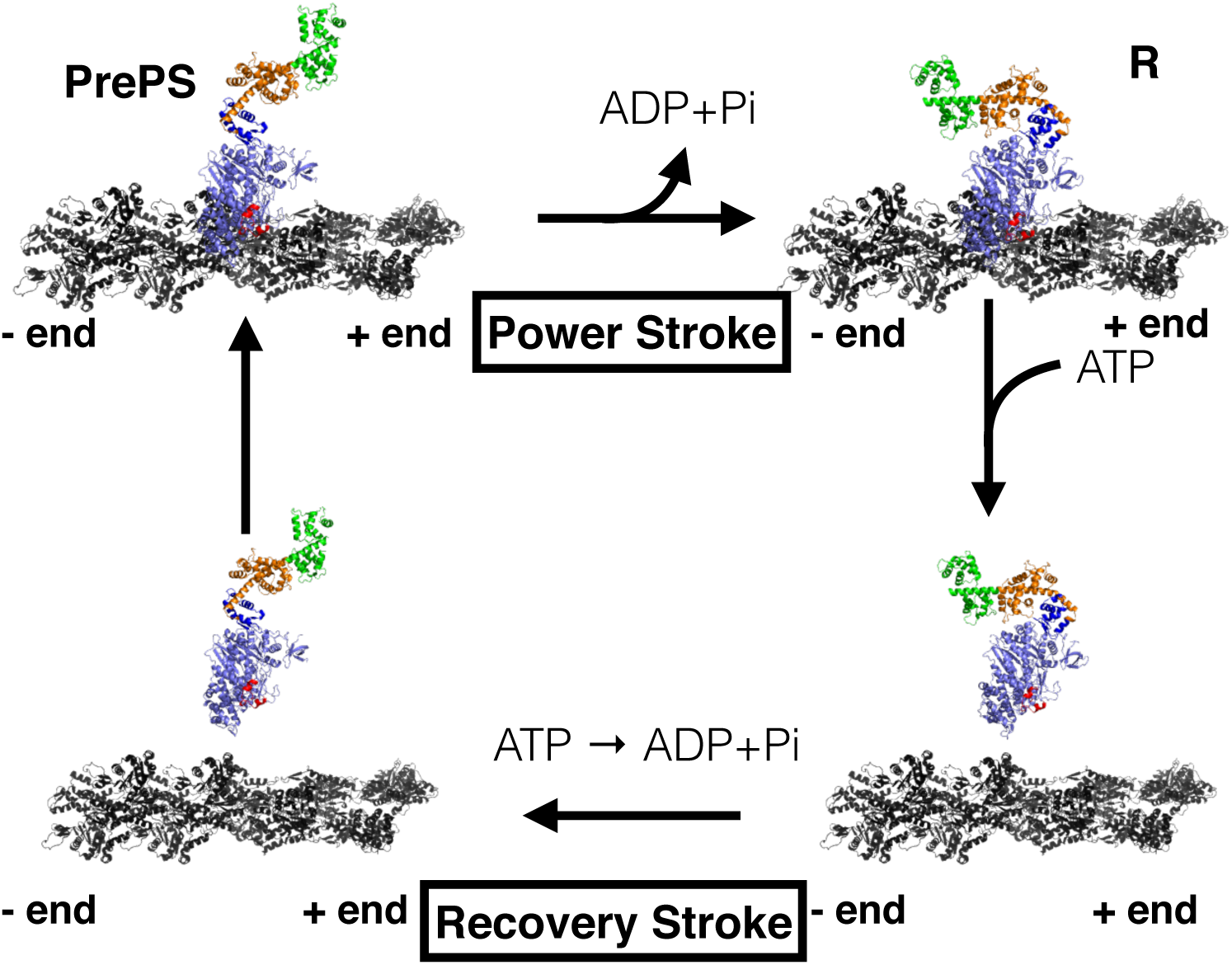
The reaction cycle of MVI. Color code. Slate blue: motor domain. Red: ins1. Dark blue: converter. Orange: ins2 and ins2-bound CaM. Green: IQ domain and IQ-bound CaM. Black: actin. The Protein Data Bank (PDB) structures used to make this figure are 4ANJ [41] and 2BKI [34] for the motor domain, MVI lever arm was extracted from 3GN4 [24], and the F-actin was adopted from 1MVW [61]. Both PrePS and R models were aligned against a myosin bound to actin in 1MVW.

Much of the work on non-muscle myosins has focused on myosin V (MV). However, since the discovery that myosin VI (MVI) has an unusual structure, there has been an increasing interest on the motility of MVI. In addition to its biophysical importance, MVI has been implicated in a wide variety of cellular functions in different organisms [3, 4]. For instance, MVI is involved in endocytosis, spermatogenesis, cell migration, in the organization of the cytoskeleton and the Golgi apparatus, and in protein localization. It also plays a role in the maintenance of stereocilia of inner ear cell. Mutations of the *myo6* gene induce deafness in mice and in humans [3]. The effect of a deafness-inducing mutation on MVI cycle has been elucidated recently, thus suggesting a mechanism by which the functioning of the mutant is hampered [5]. MVI is over-expressed in cancerous ovarian cells, and the inhibition of its expression reduces the tumor propensity to disseminate [6]. MVI is also over-expressed in cancerous prostate cells [7].

Some myosins perform their physiological function as monomers, others form dimers that walk processively, i.e. they take multiple consecutive steps on the polar track F-actin, without detaching. Although MVI is monomeric in solution [8], it dimerizes when a leucine zipper is attached distal to the lever arm [9, 10], by clustering MVI monomers on actin [11], or with the help of an appropriate protein target interacting with the C-terminal cargo binding domain of MVI [12, 13]. MVI dimers are capable of processive movement along F-actin [9, 10].

The basic principle necessary for processive movement of motors is a high duty ratio, that is the motor must spend a large part of the cycle tightly bound to actin [2]. The communication between the two heads, also known as gating, is believed to play a role in a processive dimer [2]. Kinetic models suggested that gating might not be necessary for a robust processivity if the motor displays a high duty ratio [14]. However, gating likely increases motor efficiency and run-length [14, 15]. Interestingly, recent coarse-grained models of MVI leading and trailing head provide structural evidence that the ADP release is gated [16]. These findings are in agreement with experiments [17] and kinetic models [14, 15], although some experiments suggest that blocking ATP binding to the leading head constitutes the gating mechanism of MVI [18].

Beside displaying gating and high duty ratio [19, 18], MVI shows a number of striking features that distinguish it from other processive myosins, such as the well-studied MV. (i) For starters, while MV and all other members of myosin superfamily move towards the barbed end (or + end) of F-actin, MVI steps towards the pointed end (or - end) [20, 2]. (ii) The lever arm architecture differs between MV and MVI [21]. In MV there are six calmodulins (CaMs) or light chain-bound IQ domains that constitute the lever arm, with length that is commensurate with the 36-nm spacing in the F-actin repeat. MVI has only one IQ domain, but in a step covers the same distance as MV, suggesting that other elements contribute to the lever arm, but their origin remains unclear [22, 23, 24, 25, 26, 21, 27, 28]. (iii) Furthermore, the step-size distribution of the wild-type MV is much narrower than in MVI [9, 10, 21]. (iv) Lastly, both MV and MVI move processively by a hand-over-hand mechanism [29, 30, 31], but in MVI there is evidence of inchworm-like steps [32].

Much of the unconventional mechanical properties of MVI is attributed to insert 1 and insert 2 (ins1 and ins2) (see Fig. 2), two unique fragments within the myosin superfamily [20]. Single-molecule experiments [33], and crystal structures of the R [34] and PrePS [35] states indicated ins2 as the key structural element responsible for the reversal of MVI directionality during stepping. Indeed, ins2 wraps around the converter domain, effectively turning the direction of the swing backwards, and its removal changes the directionality of MVI motion. Kinetics experiments suggest that ins1 plays a key role in determining MVI high duty ratio and gating [18, 36].

**Figure 2:**
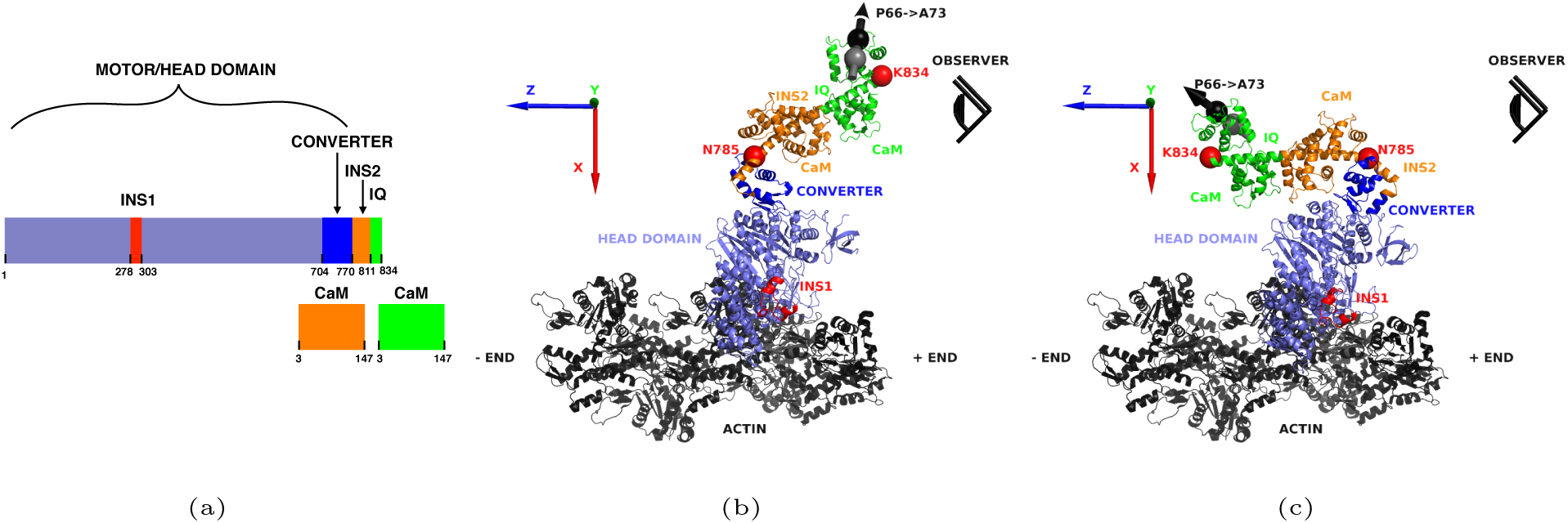
Sequence and structural models of MVI. (a) Sequence of the modeled MVI with bound CaMs. PrePS (b) and R (c) models used in the simulations. The color code is the same as in Fig. 1. The total number of residues in the protein is 834. Each CaM has 145 residues. The total number of residues for our PrePS and R models is 1124. F-actin was not part of the model, but it is shown here for reference. The red spheres show the residues N785 and K834, the beginning and the end of the lever arm in our model. The grey and black spheres show P66 and A73 of the IQ-bound CaM. The size of the four spheres is exaggerated to enhance visibility. In [39] a bifunctional rhodamine used to reveal the orientation of the lever arm is attached to these two residue. The three axis, 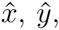 and 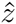 are shown as a red, green, and blue arrow, respectively. The position of the observer is shown pictorially as an eye. The observer stands on F-actin behind MVI, parallel to the 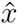 axis, and oriented in the negative direction, looking towards the pointed end of the filament.

Some of the essential features of MVI stepping mechanism are not fully understood, including the unusually large step-size distribution, and the alternation between hand-over-hand and inchworm steps. Although models of the dimer incorporating the flexibility of MVI lever arm are capable of reproducing known features of MVI motility [37], experiments using a chimeric MVI with MV lever arm showed that the step-size distribution was similar to the wild-type [33, 38], suggesting that the elusive structure of MVI lever arm might not be the key ingredient needed to explain MVI peculiar stepping mechanism. It is suspected that the putative uncoupling (or pliancy) of MVI leading-head lever arm, which was suggested by a number of experiments [39, 40, 32, 41], may contribute to the large step-size distribution. Furthermore, according to a recently proposed model [32, 38], short steps occur if the free, trailing head of a myosin dimer binds F-actin while the lever arm of the actin-bound, leading head remains in the pre-stroke orientation. Conversely, a rotation of the bound-head lever arm to the R-state orientation results in large steps.

Although experimental evidence support the model of the uncoupling of the lever arm, direct evidence requires a structural model capable of describing the dynamics of the lever arm swing. We use a combination of coarse grained (CG) simulations and theory to probe the dynamics of the swing of the lever arm in MVI. In recent years, numerous simulations on a variety of systems have shown the reliability of CG models in extracting the salient features of the dynamics of macromolecules [42, 43, 44, 45, 46]. We prepared a model for the PrePS and R conformation that include the lever arm up to ins2 and the IQ domain (Figs. 2b-c). We modeled the power stroke by inducing the transition from the PrePS to the R conformation, ignoring any intermediate configuration (see **Materials and methods** section for a fuller discussion). We ran 100 trajectories of the PrePS→R transition to study the power stroke. To understand the effect of backward load on the rotation of the lever arm, we ran 100 trajectories of the PrePS→R transition with the lever arm subject to a resistive force of 2pN, 4pN, and 6pN. Finally, we carried out 10 simulations of MVI in R state under backward load (pulling force of 1pN, 10pN, and 20pN), so to induce the reverse rotation towards the PrePS state. Our results suggest that the PrePS→R transition occurs in two major steps: in the first step, the motor domain reaches the R state, while the lever arm is uncoupled. In the second step, the lever arm diffuses towards the R state configuration with a small guidance from the motor domain. Our model leads us to propose that the inchworm-steps might occur when the free head binds F-actin before the uncoupled lever arm rotates to the R-state conformation, which identifies a direct connection between short (or inchworm-like) steps and and the uncoupling of the lever arm. The results of our simulations not only compare favorably with experiments, but also yield precise, testable predictions.

## Results

### The Lever Arm Rotation Occurs in Two Major Steps

We monitor the dynamics of the PrePS → R transition using the structural overlap function (*χ*) with respect to the R state given by [47]: 
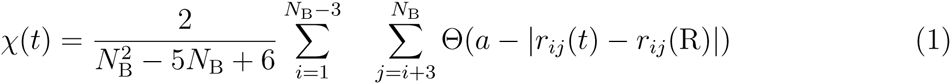
where *N*_B_ is the number of beads in the coarse grained model, *r*_*ij*_(*t*) is the distance between the beads *i* and *j* at time *t*, *r*_*ij*_(R) is the distance in the R state, Θ is the Heaviside function, and *a* = 2Å is the tolerance. The summation in Eq. 1 is over pairs of beads that are at least three residues apart in the sequence (the total number of such pairs is the inverse of the prefactor in Eq. 1). In the PrePS state, *χ ≈* 0.62, and in the R state, *χ ≈* 0.34. The time trace of *χ* shows that the PrePS*→*R transition occurs in two steps (black line in Fig. 3a). Within a few microseconds, *χ* decreases to *≈* 0.55 (inset in Fig. 3a) and it fluctuates around this value for a long time until it undergoes another rapid transition leading to the R state. The structural overlap function fluctuates around 0.34 until the end of the simulation. Similar patterns are found in all the trajectories.

**Figure 3:**
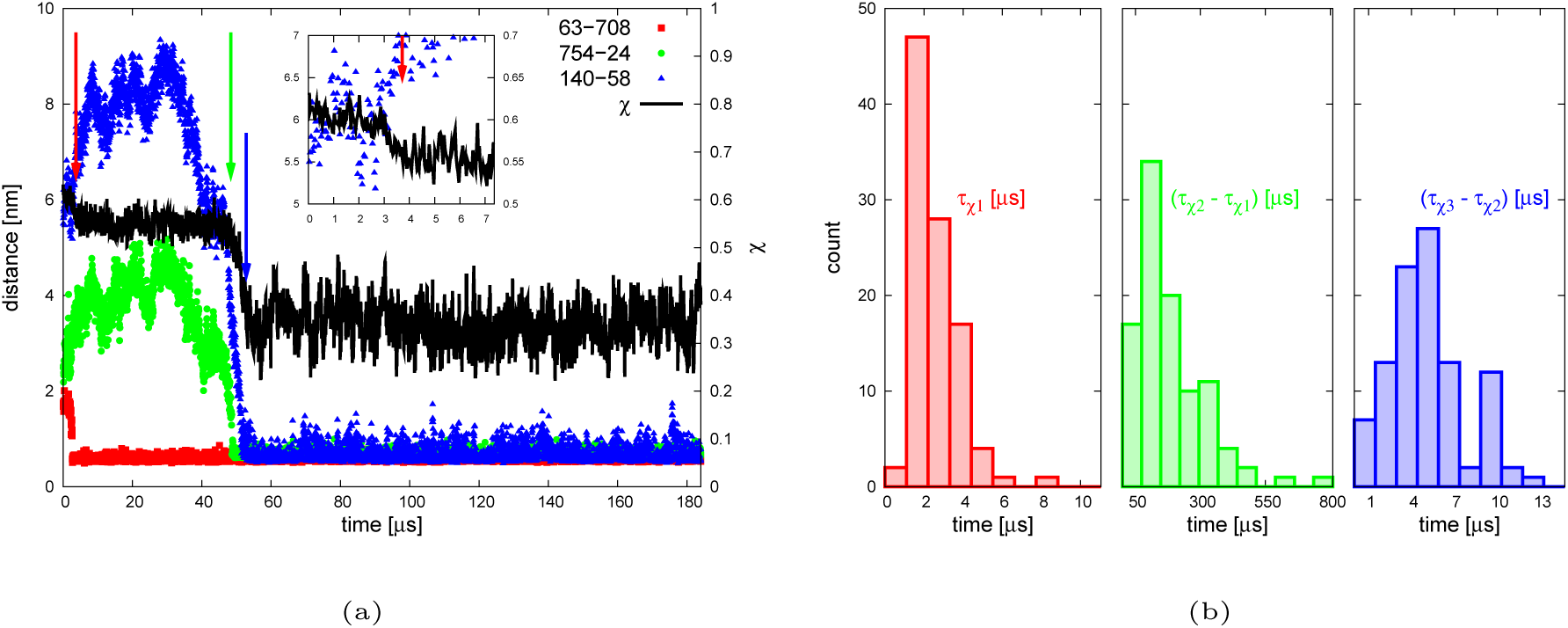
Dynamics of the PrePS→R transition. (a) The dependence of *χ* on time is in black. The red arrow indicates the end of the first transition, the green and blue arrows correspond to the start and end of the second transition, respectively. The figure also reports the time traces of the distance between C63 and R708 (red squares), T754 and D24 of the ins2-bound CaM (green circles), and V140 and D58 of the ins2-bound CaM (blue triangles). The inset is the initial transition in *χ*. (b) Left panel: histogram of *τ*_*χ*^1^_. Central panel: histogram of *τ*_*χ*^2^_ − *τ*_*χ*^1^_. Right panel: histogram of *τ*_*χ*^3^_−*τ*_*χ*^2^_. Here, *τ*_*χ*^1^_ is the end of the first transition, and *τ*_*χ*^2^_ and *τ*_*χ*^3^_ are, respectively, the start and the end of the second transition monitored by *χ*.

Let us refer to the time of the completion of the first step as *τ*_*χ*^1^_, the time to the beginning of the second step as *τ*_*χ*^2^_, and for the time to complete the second step as *τ*_*χ*^3^_. In practice, *τ*_*χ*^1^_(*τ*_*χ*^3^_) is measured as the first time *χ* reached 0.55 (0.34) (red and blue arrows in Fig. 3a). The time at which the second step begins, *τ*_*χ*^2^_, is measured as the last time the trajectory crosses 0.55 before reaching 0.34 (green arrow in Fig. 3a). The histogram of *τ*_*χ*^1^_, *τ*_*χ*^2^_.−*τ*_*χ*^1^_, and *τ*_*χ*^3^_ − *τ*_*χ*^2^_ are shown in Fig. 3b. Note the difference in the scale of the abscissa. Clearly, the first transition occurs rapidly (*τ*_*χ*^1^_ average is 〈*τ*_*χ*^1^_〉 = 2.6*μ*S), then there is a long waiting time (〈*τ*_*χ*^2^_−*τ*_*χ*^1^_〉 = 181 *μ*S) before another fast transition (〈*τ*_*χ*^3^_−*τ*_*χ*^2^_〉 = 5.1*μ*S).

### Structural Transitions During the Two Steps

In order to characterize the structural origin of the two steps observed in the time trace of *χ*(*t*) during the PrePS→R transition, we monitor the following three distances between beads that form native contacts in the R state, but not in the PrePS state: (i) dRED is the distance between C63 and R708 (red spheres in Fig. 4), (ii) the distance between T754 and D24 of the ins2-bound CaM is dGREEN(green spheres in Fig. 4), (iii) dBLUE is the distance between V140 and D58 of the ins2bound CaM (blue spheres in Fig. 4). When the converter moves from the PrePS to the R-state conformation, the dRED contact is formed. The formation of the dGREEN contact indicates the initial closure of the lever arm from its open, PrePS orientation, to the locked position of the R state, and dBLUE permits us to monitor the final closure of the lever arm onto the motor domain. The time traces of these three distances for a particular trajectory are shown in Fig. 3a, from which it appears that the formation of the dRED contact is simultaneous to the first step of *χ*(*t*), and the second step of *χ*(*t*) occurs around the same time when dGREEN and dBLUE are formed. To determine whether this observation holds for all trajectories, we extract from each trajectory the time for stable formation of the three contacts. By “stable” we mean that once the contact is formed, it remains intact with small fluctuations (in practice, we monitor stable contacts by checking the first time at which the contact is formed such that for the rest of the simulation the contact stays closer to the R-state ideal distance than to the largest distance explored during the PrePS→R transition). We refer to the time for stable formation of dRED, dGREEN, and dBLUE as *t*_1_, *t*_2_, and *t*_3_, respectively, and we investigate the correlations between *t*_1_ and *τ*_*χ*^1^_, *t*_2_ and *τ*_*χ*^2^_ and *t*_3_ and *τ*_*χ*^3^_. Figure 5a shows that there is good correlation between *t*_1_ and *τ*_*χ*^1^_, with the exception of a few points marked as filled, which represent all those trajectories in which the lever arm begins to close onto the motor domain before the rotation of the converter domain occurs, that is before dRED is formed. In these cases, the initial drop in *χ*(*t*) reflects the formation of the dGREEN contact. In three out of five cases, the initial dGREEN contact is transitory, which means that it rapidly breaks, the converter rotates and it finally reforms, thus ultimately the stable formation of the dRED contact precedes the dGREEN contact. In the two cases highlighted with an arrow in Fig. 5a (see also Fig. S19) the dGREEN contact is stably formed before dRED, and it does not break. The correlation coefficient and the linear fit, performed in the absence of the filled points, illustrates that in most cases (95% of the trajectories) dRED forms around *τ*_*χ*^1^_. This suggests that at *τ*_*χ*^1^_ the converter is on the +end side of MVI motor domain (see Figs. 2b-c). A close examination of the crystal structures before and after the PrePS→R transition [34, 35] reveals that the converter undergoes also a rotation and it is subject to a conformational change. The distance dRED can only monitor the translation of the converter, but we show in the SI that around *τ*_*χ*^1^_. the converter undergoes the rotation and the structural transition necessary to reach the R-state conformation. We deduce that, in most of the cases, the first step in the lever arm swing corresponds to the movement of the converter from the PrePS to the R configuration.

**Figure 4:**
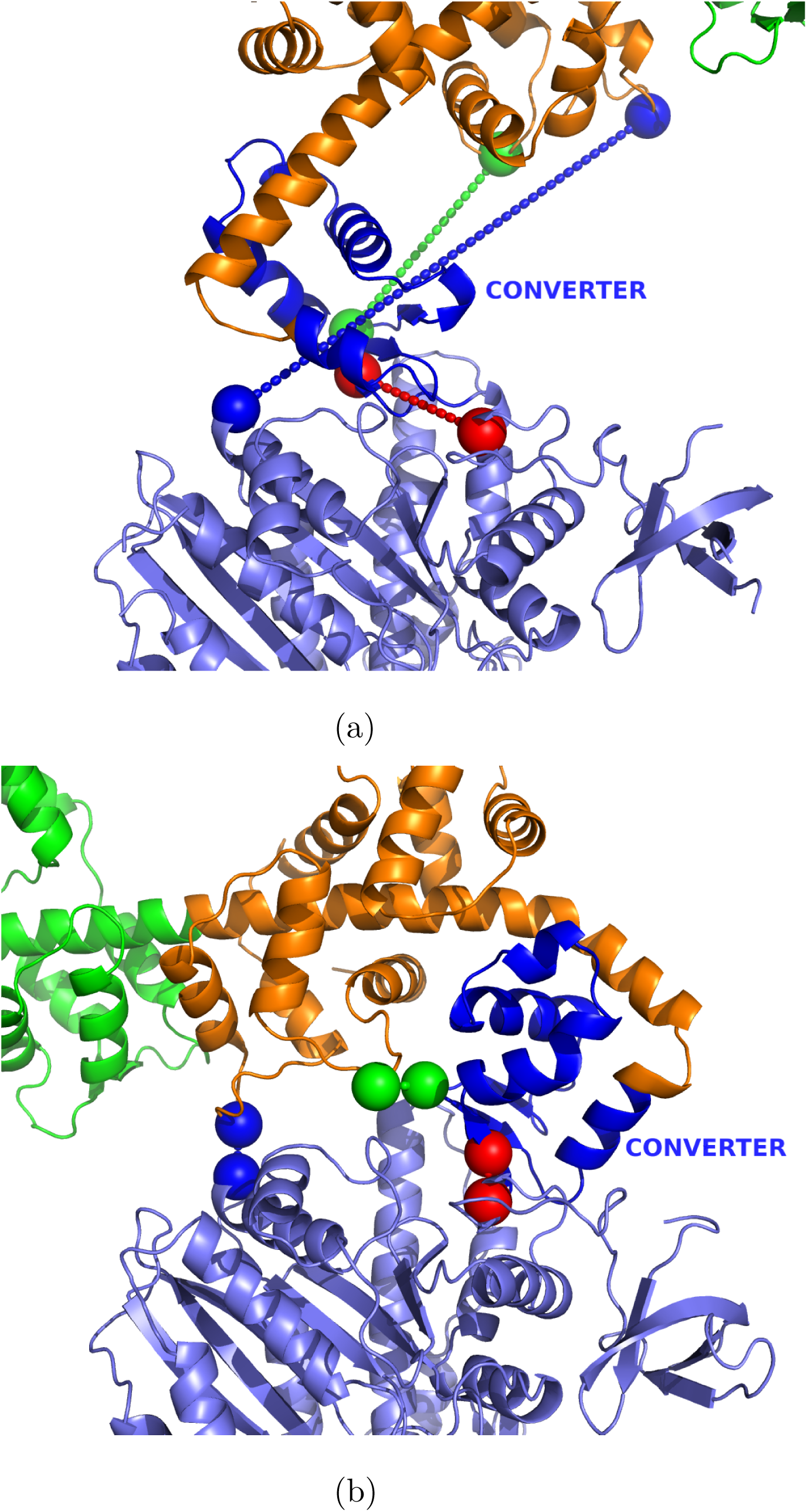
Monitoring the key structural changes in the PrePS R transition. The locations of the C_*α*_ atoms of C63 and R708 are shown as red spheres, and those of T754 and D24 of the ins2-bound CaM are shown as green spheres. The blue spheres show the C_*α*_ positions of V140 and D58 of the ins2-bound CaM, respectively. The dashed lines connect the beads involved in the dRED, dGREEN, and dBLUE contacts. The PrePS state is in (a), the R state is in (b).

**Figure 5:**
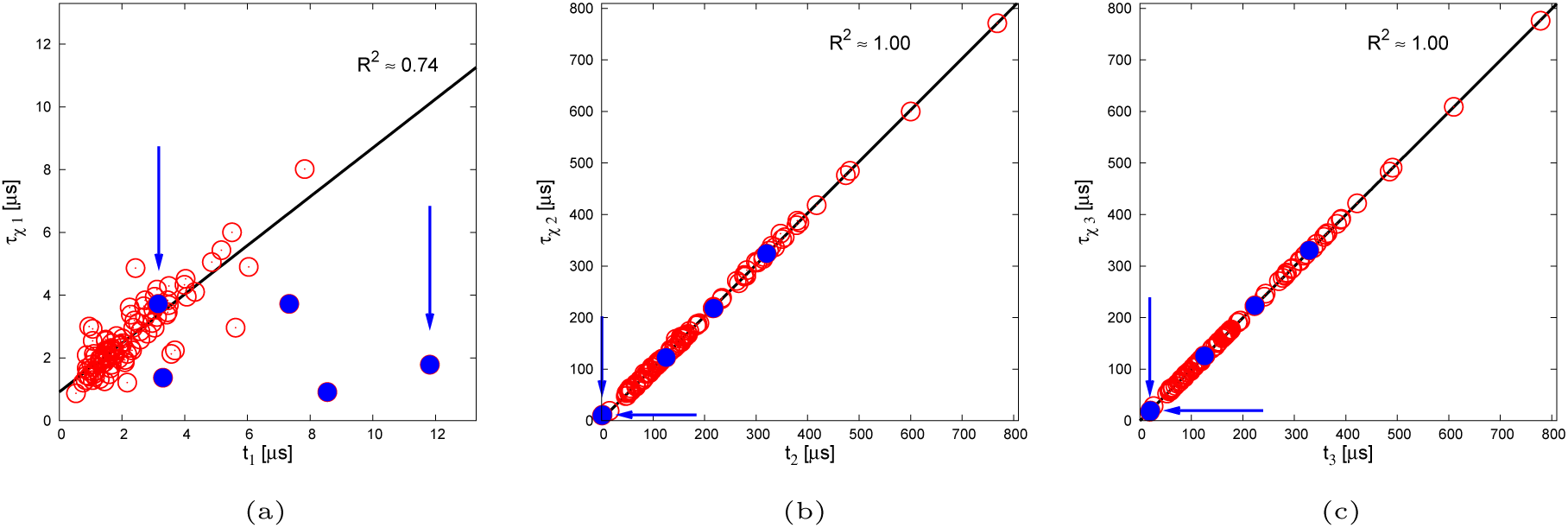
Correlation between transition in *χ* and the structural changes in MVI. (a) Plot of *τ*_*χ*^1^_ against *t*_1_. Blue circles indicate results from simulations in which temporarily (or permanently, if indicated with an arrow) the dGREEN contact forms prior to the dRED contact. The slope of the linear fit is ≈ 0.78, while the intercept is ≈ 0.89*µ*s.(b) Plot of *τ*_*χ*^2^_ versus *t*_2_. The slope of the linear fit is ≈ 1.00, and the intercept ≈ 2.8*µ*s. (c) *τ*_*χ*^3^_ plotted against *t*_3_. The slope of the linear fit is ≈ 1.00, and the intercept ≈ − 0.20*µ*s. The linear fits and the correlation coefficients are computed without the blue-filled points in (a), and with all the points in (b–c).

The correlations of *t*_2_ with *τ*_*χ*^2^_ and of *t*_3_ with *τ*_*χ*^3^_ are shown in Figs. 5b-c. The two correlation coefficients (in this case computed accounting using all the points) are very close to unity, and the linear fit yields small intercepts (see caption in Fig. 5). Hence, the dGREEN contact forms around *τ*_*χ*^2^_ and it is followed by the formation of the dBLUE contact at about *τ*_*χ*^3^_.

The analysis conducted so far monitoring *χ* and the formation of the dRED, dGREEN, and dBLUE contacts suggest that we can divide our trajectories in two classes: (1) In most cases (95%), the first step of *χ* corresponds to a movement of the converter domain from the PrePS to R position (formation of the dRED contact). Rarely (3%), we observed that the movement of the converter is preceded by the transitory formation of a R-state contact between the lever arm and the motor domain (dGREEN contact). Once the converter has moved, after a long waiting time, ins2 closes onto the motor domain to form the R state (stable formation of dGREEN and dBLUE contacts). Hence, the first step is a power stroke transition of the motor domain, and during the second step the lever arm attains the R-state conformation. This suggests that, in all these trajectories (98 out of 100), the movement of the lever arm is “uncoupled” from the PrePS→R transition of the converter. We refer to these trajectories as “uncoupled”. An example of an uncoupled trajectory is shown in Movie1 and Movie2. (2) In a minority of trajectories (2%) a R-state-like stable contact between ins2 and the motor domain is formed before the switch of the position of the converter. These trajectories are classified as “coupled”, as the interaction between the lever arm and the motor domain leads to a rapid swing of the lever arm directed by the movement of the converter. Movie3 and Movie4 show an example of a coupled trajectory.

Because coupled trajectories are only rarely observed with or without backward load (see SI text and Fig. S19), we focus the rest of our analysis on the uncoupled trajectories, relegating a fuller discussion of the coupled trajectories to the SI. In particular, we want to quantify the extent of the uncoupling of the ins2-IQ region of MVI after *τ*_*χ*^1^_, and devise a simplified model to understand its dynamics.

### The Rotation of the Ins2-IQ domain in the Uncoupled Trajectorie

The reference system is shown in Figs. 2b-c. The 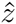 axis is nearly parallel to the actin filament, with the -end of F-actin directed towards the positive side of the axis. MVI is approximately parallel to the 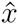 axis, and oriented towards the negative direction. The view of the observer used as reference is described in Figs. 2b-c. To follow the movement of the lever arm, we monitor the position of N785 (in ins2, shown as a red bead in Figs. 2b-c) and K834 (last residues of the IQ domain, shown as a red bead in Figs. 2b-c). A sample trajectory (Fig. S5, Movie1, and Movie2) shows that N785 moves during *t* < *τ*_*χ*^1^_ from PrePS-state nearly to the R-state configuration, and when *t* exceeds *τ*_*χ*^1^_ it rapidly finds the R-state equilibrium position, fluctuating around this value for the rest of the simulation. On the other hand, K834 undergoes a large rotary movement from the value in the PrePS state to the one in the R state (Fig. S5, Movie1, and Movie2). We conclude that, after the initial step in the PrePS→R transition, N785 serves as the hinge around which the distal part of ins2 and the IQ domain rotate until they reach the R state. Let the segment of sequence N785-K834 be the lever arm. To monitor the rotation of the lever arm in the most intuitive way, we describe the vector connecting N785 to K834 in spherical coordinates. Since the distance between N785 and K834 is roughly constant during the PrePS→R transition (top panel in Fig. S6), we describe the movement to the R state of the uncoupled lever arm as a rigid rod rotation with changes in the altitudinal angle, *θ*, and the azimuthal angle, *φ* (Fig. 6a).

**Figure 6:**
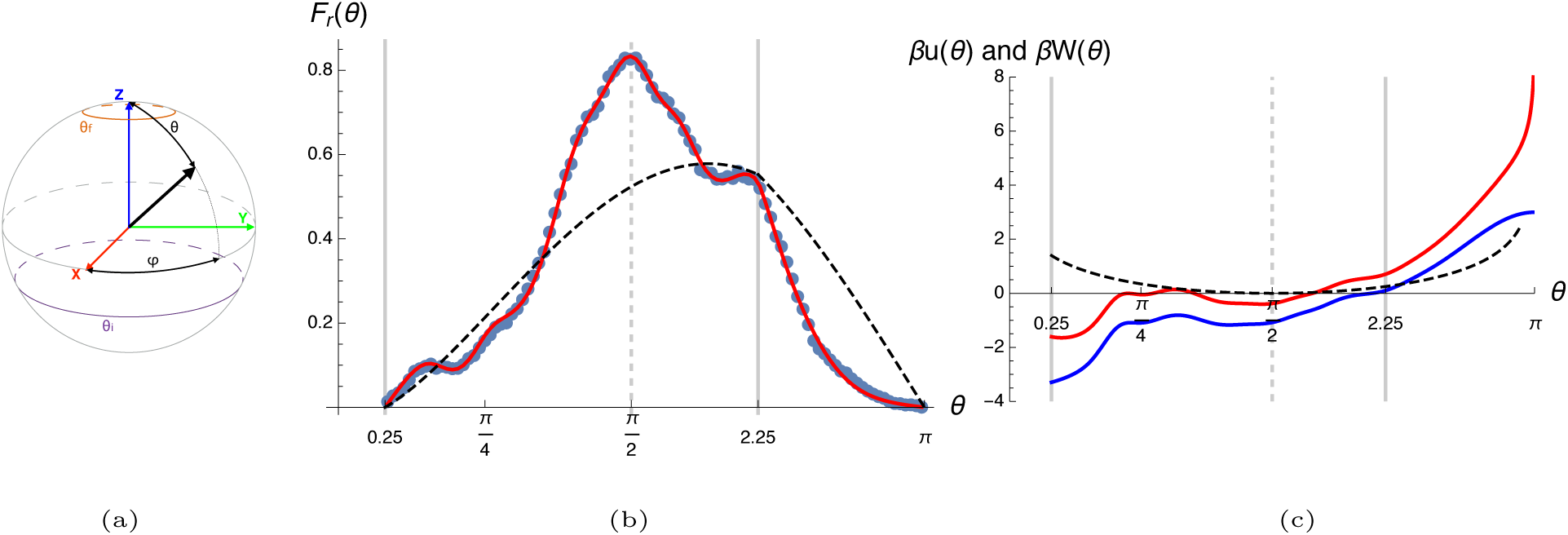
Extraction of the potential driving the rotation. (a) Definition of the the angles *θ* and *φ*. The cartesian axes are shown in red, green, and blue. The direction of the lever arm is shown as a black arrow. Note from Fig. 2b-c that the z axis is parallel to F-actin, so the *θ* angle indicates the angle between the lever arm and the actin filament. The source and sink of the FP model in Eq. S20 are approximately shown in purple and orange, respectively.(b)The probability *F*_*r*_(*θ*). The result from coarse grained simulation is shown in blue dots. The probability distribution obtained for the case of free rotation (*u*(*θ*) = 0 and *σ*_*φ*_(*θ*) = 1) is shown as black, dashed lines. The fit of the data, obtained using Eq. S29, is in red. The grey thick lines show the values of *θ*_*i*_ = 2.25 and *θ*_*f*_ = 0.25. (c) The potential obtained to fit *F*_*r*_(*θ*). The red line is *W*(*θ*) = *u*(*θ*) *k*_*B*_*T* log [sin(*θ*)*σ*_*φ*_(*θ*) (where *σ*_*φ*_(*θ*)^2^ is the variance of the distribution in *φ*, see Eq. S30, Figs. S12-S13 and discussion in SI), and the blue line is *u*(*θ*). The grey thick lines are the same as in (a). The black dashed line shows *βW*(*θ*) for a free rotator.

### Energetics of the Lever Arm Swing

The rotating lever arm is subject to a potential *U* (*θ, φ*) that governs its motion. We want to extract this potential from the trajectories to decipher the extent of the coupling between the lever arm and the motor domain. Assuming that at the start of the rotation the lever arm is at an angle *θ* = *θ*_*i*_, at the end the altitudinal angle is *θ* = *θ*_*f*_x, and ignoring the role of the azimuthal angle, a large (*U*(*θ*_*i*_, *φ*′) − *U*(*θ*_*f*_,*φ*)>>*k*_*B*_*T*), downhill potential suggests that the swing is guided. In contrast, if the potential is flat (*U* (*θ*_*i*_,*φ*′) − *U*(*θ*_*f*_,*φ*)*≈k*_*B*_*T*), then the lever arm is uncoupled in the *θ* direction. To extract the potential *U*(*θ*, *φ*), we cannot simply compute the logarithm of the equilibrium probability distribution *ρ*(*θ*, *φ*), because once the lever arm reaches the R-state we never observe a fluctuation to the PrePS-state. (In fact, we found out that even a rearward load of 10pN is not sufficient to generate such backward rotation during a 368*µ*s-long simulation, see SI text and Fig. S18). Hence, the system is out-of-equilibrium, and we need to adopt a different strategy to extract *U* (*θ*, *φ*). We adapt a procedure put forward in [48] to our case (see details in SI). We generate a stationary probability distribution on a unit sphere *ρ*_*r*_(*θ*, *φ*) from the simulated PrePS→R transitions. The trajectories are injected at an angle *θ*_*i*_ = 2.25rad(close to the PrePS state, *θ*_PrePS_ *≈* 2.55rad), and are removed when they cross *θ*_*f*_ = 0.25rad (close to the R state, *θ*_R_ *≈* 0.04rad). We assume that the stationary probability *ρ*_*r*_(*θ*, *φ*) is the solution of a Fokker-Planck (FP) equation for a rotating rod subject to a potential *U* (*θ*, *φ*) [49], and equipped with the appropriate boundary conditions to account for injection and removal of the trajectories [48]. Since we know *ρ*_*r*_(*θ*, *φ*) from simulations (see SI and Figs. S10a,c), we obtain the potential *U* (*θ, φ*) by solving the FP equation. To solve the FP equation we assume that: (i) the diffusion coefficient *D* is a scalar constant, and (ii) the *φ*-dependence of the potential *U* (*θ*, *φ*) is harmonic with a *θ*-dependent spring stiffness, which we extract from the trajectories assuming a Boltzmann distribution in the azimuthal angle. These two assumptions are approximations: in general, the diffusion coefficient is a space-dependent tensor, and since the time scale of the dynamics in *θ* and in *φ* is roughly the same (*∝* 1/*D*), there is no guarantee that the azimuthal angles obey a Boltzmann-like distribution. Using these approximations, we obtain a one-dimensional FP equation in *θ*, whose solution Eq. S29 connects the probability distribution 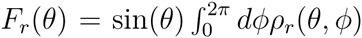 to a free energy *W*(*θ*) (Eq. S30), which has an energetic term arising from *U*(*θ*, *φ*) (which we refer to as *u*(*θ*)), and an entropic contribution resulting from the dimensionality reduction (from two dimensions, *θ* and *φ*, to only *θ*). In Fig. 6b we show the probability distribution 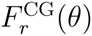 obtained from the analysis of the coarse-grained simulations (blue dots). As described in detail in the SI, we fit the free energy *W*(*θ*) to obtain a probability distribution *F*_*r*_(*θ*) (red line) that closely recovers the results from simulation. As a comparison, we also show in Fig. 6b as a black, dashed line the probability distribution in the case of free rotation.

The profile of *W*(*θ*) (Fig. 6c, red line, the blue line is *u*(*θ*)) shows that a conformation in which the lever arm points backwards (*θ ≈ π*) is unlikely for both entropic and energetic reasons. From *θ* = 2.25rad (*≈* 129°) to *θ ≈ π/*4 (*≈* 45°) a large part of the swing occurs on an almost flat energy profile. For *θ < π/*4 the energetic term of *W*(*θ*) (*u*(*θ*)) contributes with a shallow minimum of *≈*1 kcal/mol, that drives the last part of the swing. The free energy profile for the freely rotating lever arm is shown as a black dashed line in Fig. 6c. We conclude that the swing occurs without a strong guidance of the motor domain, implying that the lever arm movement is mostly uncoupled, which is qualitatively revealed in the trajectories (see Fig. S8, Movie1, and Movie2). This means that for large part of the swing, the lever arm rotates stochastically while maintaining a hinge around N785. The capture towards the R state occurs only when the lever arm is sufficiently close to the the motor domain.

To assess the validity of the approximations used to solve the FP equation, we compare the simulated mean first passage time (MFPT) 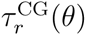 and the one resulting from a diffusing pseudo-particle in the potential *W* (*θ*). The theoretical MFPT is obtained by solving the one-dimensional FP equation using a constant diffusion coefficient [50],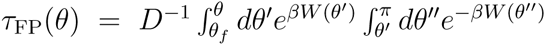.Fitting *D* in *τ*_FP_(*θ*) versus 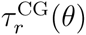 we obtain *D≈* 0.01/*µ*s(see SI). The good agreement between *τ*_FP_(*θ*) and 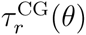 reported in Fig. S15 suggests that the assumptions made in solving the FP equation are reasonable. Given the value of the diffusion constant *D*, a freely rotating lever arm (*βW*(*θ*) = *−* log[sin(*θ*)], the black dashed line in Fig. 6c) undergoes the rotation *θ*_*i*_ → *θ*_*f*_ in *≈* 396*µs*, which is slower than the mean first passage time obtained from CG simulations (*≈* 176*µs*). To estimate the extent of guidance of the motor domain, we define a potential linear in *θ*, *u*(*θ*) = *k*(*θ − θ*_*i*_), and we extract the value of *k* such that the *θ*_*i*_ → *θ*_*f*_ rotation occurs in the same amount of time as observed in the CG simulations. We recover a MFPT of *≈* 176*µs* with *k*(*θ*_*i*_ − *θ*_*f*_) *≈* 2.05*k*_*B*_*T*, which again suggests that the rotation of the lever arm is subject to a small drift. Thus, it is physically reasonable to propose that the coupling of the swing to the motor domain is weak.

Our approximations yielded a one-dimensional profile that captures the features of the CG simulations. However, the spatial dependence of the diffusion tensor, and the non-equilibrium nature of the dynamics in the *φ* direction are likely to play a role in the 2D probability distribution and MFPT, as shown in Figs.S10a-d.

### Comparison With Experiments

We use our simulations to compare with experiments probing the stepping mechanism of MVI dimers [39]. To follow the orientation of the lever arm during the stepping, the CaM bound to the IQ domain of MVI was labelled with a bifunctional rhodamine (BR) probe. The points of attachment are P66 and A73, and are shown in Figs. 2b-c in black and grey, respectively. Sun *et al.* inferred the rotational dynamics of the lever arm from the orientational changes in BR using polarized Total Internal Reflection Fluorescence (polTIRF). This yielded the *θ* angles (along the direction of actin, referred as *β* in [39]) and the *φ* angle (the azimuthal angle is named *α* in [39]). (Note that *θ* and *φ* are angles of the BR probe with respect to the reference system of actin (Figs. 2b-c). In [39], the greek letters *θ* and *φ* are used to identify the angle of the lever arm in the reference system of actin, and *α* and *β* are used for the probe. We compare here with *α* and *β* of [39].) The head of the dimer in the leading and in the trailing positions were identified, and the corresponding distributions of the *θ* and *φ* angles were extracted. According to kinetics experiments [51],both the trailing head (TH) and the leading head (LH) are in the ADP-bound state. In the TH, the lever arm swing has already occurred, and it is directed towards the pointed end of F-actin. In contrast, the LH, held under tension by the actin-bound TH, is in the uncoupled state (from now on referred to as U).

To compare our simulations with experiments, we reproduced the measurements by monitoring the orientation of the unit vector connecting A73 with P66 (Figs. 2b-c). The *θ* and *φ* angles of the BR probe in the R-state crystal structure differ between the computational model (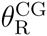 and 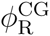) and the values used in [39] (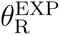 and 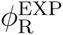). Two factors likely contribute to this discrepancy: (i) the alignment with F-actin and the choice of the reference system is not identical in experiments and simulations (our axes are rotated compared to [39], so *φ ≈ α* + *π/*2), and (ii) we extracted the angles from the relative position of the *α*-carbons of P66 and A73 from CaM, while experiments detect the orientation of the *β*-carbons [52]. Thus, we correct for this error by removing from all the probe angles *θ* and *φ* extracted from simulation the quantities 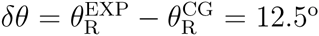 and 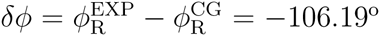 respectively.

To describe the TH, we consider the R state conformations sampled in the simulations only at *t* > *τ*_*χ*^3^_, to ensure that the R state is reached. To sample the U state of the leading head, we ran 100 simulations of the swing of the lever arm by applying to the tip of the lever arm (K834) a backward load of 6pN (details in SI), which mimics the effect of the inter-head tension. The mechanical force (*f*_*L*_) adopted to describe the U state resists the rotation of the lever arm by applying a momentum of the force of magnitude *≈r*_*LA*_*f*_*L*_, where we consider a lever arm *r*_*LA*_ *≈*7.5nm (Fig. S6, top panel). If we consider the entire lever arm of MVI (*r*_*LA*_ *≈*18nm), the force that generates the same resistive momentum is *f*_*L*_ *≈* 2.5pN, which is close to the upper bound of the inter-head tension estimated experimentally (*f*_*L*_ *≈* 2.2pN, [17]). This suggests that the magnitude of our backward load is not unrealistically high. We show in the SI that the backward load of 6pN does not affect the first step of the PrePS→R transition (see Figs. S2-S3, SI text, Movie 5, and Movie 6), but it is sufficient to prevent the occurrence of the rotation forward of the lever arm (second step, see Movie 5 and Movie 6). Thus, we generated the ensemble of U configurations as the collection of structures from the simulations conducted with 6pN backward load, sampling only the configurations for *tτ*_*χ*^1^_.

Comparison of experimental data and simulations is provided in Fig. 7. We compared the following quantities reported in [39]: (i) the probability distribution for the LH and TH of the *θ*-angle (Fig. 7a), (ii) the probability distribution for the change in the *φ*-angle after one step (∆*φ*, Fig. 7b), (iii) the probability distribution for the change after two steps of both the *θ*angle (^2^∆*θ*, Fig. 7c), and (iv) the *φ*-angle (^2^∆*φ*, Fig. 7d). Because in experiments there is no control on the landing azimuthal angle of MVI on actin, the experimental distribution of the *φ*-angle for the LH and TH is almost flat across 180° (Y. Goldman, personal communication, June, 2016). In simulations, we always start from a MVI parallel to the x-axis, thus the comparison of calculated and measured distributions could be misleading (see SI text and Fig. S20).

**Figure 7:**
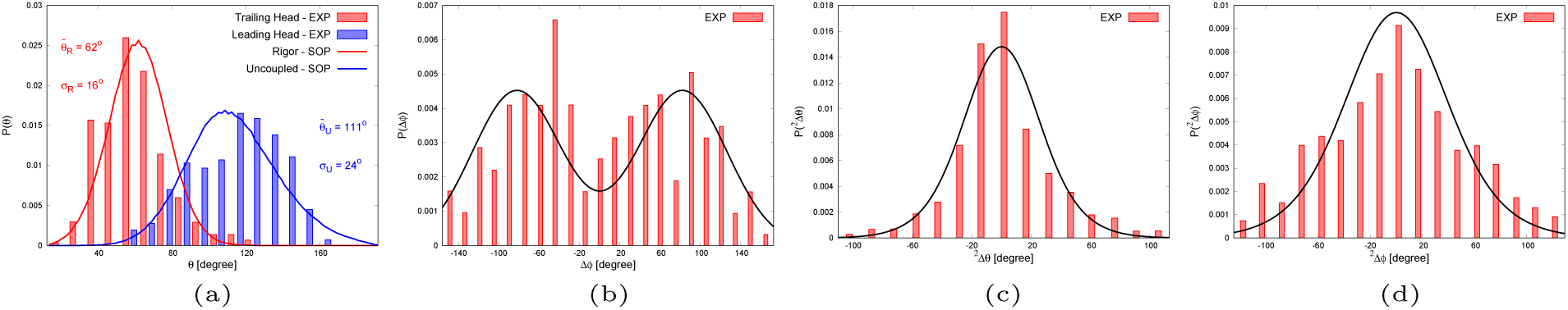
Comparing simulations and experimental results. In all the panels the histograms show the data extracted from Figs. 3-4 of [39], while the lines show the results of our simulations and analyses. The red and blue lines are obtained directly from simulations, the black lines from our model of a straight step of MVI dimer (see SI text and Eqs. S37-S38, based upon the fit of the simulated distributions in panel (a) and in Fig. S20. (a) Distribution of *θ* (*β* in [39]) of the trailing (red) and leading (blue) heads. To match the figures we displaced our data by 12.5°, which is the difference in the angle reported in [39] after aligning the crystal structure of the R state to the actin filament, and the same angle obtained after our alignment to the actin filament. The data from simulations were fit using normal distributions. The fit parameters are shown in the figure. The average of the distribution is denoted with a bar, the standard deviation is labeled as *σ*. (b) Distribution of the change of *φ* after one step (Eq. S37). (c) Distribution of the change of *θ* after two steps (Eq. S38). (d) Distribution of the change of *φ* after two steps (Eq. S38).

For comparison (i), we extracted directly the probability distributions from the CG simulations of the U state and the R state to describe the LH and TH, respectively. We did not simulate consecutive steps. Thus, for the comparisons (ii-iv) described above we created a simple model of the changes in the angles after one or two steps using two assumptions (details in SI): (1) the dimer always steps by a hand-over-hand mechanism, and (2) the values of the angles after a step do not depend on the values before the step.

We found a remarkable agreement between the *θ*-angle obtained is simulation and experiment for both the LH and TH (Fig. 7a). Our CG model was clearly able to reproduce the orientation and the fluctuations of the BR probe in the LH and TH, which validates the model. It should be stressed that we did not adjust any parameter in the CG model to obtain agreement with experiments.

Although the calculated changes in the *φ*-angle after one step agree with of the experimental data (Fig. 7b), there is a difference in the interpretation of the results based on simulations and experiments. According to experiments, in both the transitions LH→TH and TH→LH,∆*φ* can be positive or negative with roughly equal probability (see Table 1 in [39]). From simulations we find that ∆*φ* is mostly (about *≈*97% of the time) positive (negative) in the R→U (U→R) transitions. Thus, in our simulations each peak of the distribution in Fig. 7c draws contribution from either R→U or U→R transitions, whereas in experiments each transition contributes to the two peaks almost equally. In our model of stepping we assume that the direction of the inter-head tension is always aligned with F-actin, which implies that only a 36-nm steps occur. But the measured broad step size distribution in MVI suggests that this is not always the case, thus it is not surprising that our simple model of stepping does quantitatively reproduce some of the features observed in experiments.

Comparison of the probability distribution after two steps yields accurate results for the *θ*-angle (Fig. 7c), while our model for the distribution of ^2^∆*φ* results in a somewhat more peaked distribution (Fig. 7d). This suggests that, in our model of stepping, MVI is not as “wiggly” as inferred from experiments [39]. A more quantitative comparison with experiments requires a coarse-grained model of MVI dimer in complex with F-actin.

## Discussion

### Position of the hinge of the rotatory movement

We suggest that the location of the hinge around which the lever arm rotates is close to residue N785, at the beginning of a region that has already been identified as pliant in the PrePS state [41]. This residue is in ins2, within a few microseconds from the start of the PrePS→R transition it binds the converter, and then fluctuates around its average position (Fig. S5). Experiments have shown that chimeric constructs in which ins2 was truncated before (after) N785 and replaced with an artificial lever arm are + ended (− ended) motors [53, 14]. Thus, from the view point of the architecture the location of the hinge that we suggest is physically reasonable.

### The Uncoupling of the lever arm during power stroke is intrinsic to the motor head

The occurrence of a two-step PrePS→R transition with uncoupling of the lever arm at zero backward load suggests that it is not the presence of an actin-bound trailing head that induces the uncoupling of the lever arm. Our simulations illustrate that there is a possibility of a coupled swing to occur. However, this is unlikely at zero force, and was observed only once (at 4pN) whenever a backward load was applied to the tip of the lever arm (see Fig. S19). Hence, the mechanism of uncoupling is inherent in the power stroke of MVI, and it is likely not due to gating or rearward tension.

### The two-step mechanism

It has already been suggested that the power stroke of MVI occurs with a two-step mechanism, in which first the converter rotates, and then, once the backward load from the actin-bound trailing head is relieved, the lever arm rotates towards the R-state configuration [41]. In this previous model, it was argued that at the end of the first step the converter domain is in a PrePS state-like configuration, but it has moved towards the R-state position on the motor domain. This was rationalized by speculating that a transition to the R-state conformation of the converter would have resulted in a large torque. We observe that the converter domain not only translates and rotates to the R-state position on the motor domain (see SI text and Fig. S2), but it also assumes the conformation observed in the R-state crystal structure (the so-called, R-fold of the converter, see Fig. S3). The flexibility around the hinge allows for this changes to occur even in the presence of a large backward load (Figs. S2-S3 and SI text). It is worth noting that we are not considering the effects of an azimuthal component of the backward load, which might add more complexity to the picture, as we suggested in our comparison to the experimental results.

### The lever arm of MVI undergoes a counterclockwise rotation

Both the coupled and uncoupled trajectories suggest that the structure of the converter domain and the PrePS→R transition favor a counterclockwise rotation of MVI lever arm, as seen by the observer in Fig. 2b-c (see also Movie2 and Movie4). This suggests that in a stepping dimer the trailing, free head swings preferentially over F-actin. As already pointed out in [39], this might favor the binding of the free head to F-actin monomers 6 and 15 (counting from the monomer to which the leading head is bound), which would confer MVI the experimentally observed right-handed twirling around the actin filament [39, 21].

### The connection between the uncoupled state and the occurrence of short (inchworm-like) steps

A model was proposed to explain the presence of short inchworm-like steps in MVI motility [32]. The authors suggest that a short step occurs if the free MVI head binds actin while the lever arm of the leading, actin-bound head is still in PrePS orientation, whereas if the actin-bound MVI lever arm rotates to the R-state orientation, the dimer takes a long step. This scheme provides a structural picture that encompasses both the long and short steps of MVI, thus explaining the broad step-size distribution. Based on our two-step mechanism for MVI power stroke, we propose that the short (inchworm-like) steps occur if the trailing head rebinds F-actin before the rotation of the uncoupled lever arm occurs, that is before the completion of the second step of the PrePS→R transition. If the second step of the transition is completed, then the dimer takes a long step. Because ATP hydrolysis and the recovery stroke occur before the rebinding of the free head to F-actin, for our proposition to be valid the rotation of the uncoupled lever arm must occur on a timescale comparable with the rate of ATP hydrolysis, which was estimated to be *≈* 10s^*−1*^ *−* 17s^*−1*^ [54]. Our predicted timescale of transition from the uncoupled state to the R state is 〈*τ*_*χ*^3^_ − *τ*_*χ*^1^_〉 ≈ 186*µ*s, which is obtained for a lever arm of about 7.5nm. The large steps of MVI dimers demand a combination of traditional lever arm and lever arm extension of roughly 36nm for the dimer. If we assume that the timescale of rotation scales with the length of the rotating rod as *L*^3^*/*ln(*L/*2*a*) [49], where *a* is the thickness of the rod (considering CaM it is approximately 3nm), the resulting timescale becomes about 2.6ms, which is only *≈* 20 times faster than the highest bound for the measured rate of ATP hydrolysis. This suggests that it is plausible that the trailing head might bind before the lever arm of the leading head reaches the R state (that is after *τ*_*χ*^1^_, but before *τ*_*χ*^3^_).

### Uncoupling of the lever arm and phosphate release

In MV it was shown that, if the motor is pulled backward at high forces (*>*2pN), the phosphate (Pi)-release pathway opens allowing for rebinding of Pi, and a high concentration of Pi increases the rate of detachment [55]. Because Pi-release appears to precede the movement of the converter domain [56], and the converter is mechanically connected to the nucleotide binding domain through the Relay helix, we surmise that in MVI the closure of the Pi-release gateway occurs before the uncoupled lever arm swings, and it might be concurrent with *τ*_*χ*^1^_, when the converter reaches the R-state conformation. Since we do not see a significant dependence of *τ*_*χ*^1^_ on backward load, we predict that, in contrast to MV, the concentration of Pi should not affect the rate of detachment from actin of a MVI head under backward load. Because during the PrePS→R transition the converter moves backward, if the Pi-release pathway closes around *τ*_*χ*^1^_, it is hard to imagine that larger backward loads would move the converter forward and open the Pi-release gateway again.

### Predictions

Our simulations lead to the following predictions. (i) The swing of MVI lever arm occurs in two steps: first the converter reaches the R-state conformation while the lever arm points towards the barbed-end of F-actin and is uncoupled from the motor, then the uncoupled lever arm rotates until it reaches the R-state conformation. The first step is only mildly dependent on backward load. The uncoupling of the lever arm from the motor domain after the first step suggest that, in contrast to MV [55], the excess of Pi in solution should not lead to a faster detachment from F-actin of a backward-pulled MVI. (ii) The hinge around which the lever arm rotates is close to N785. (iii) The energetics of the lever arm during the rotation suggests that it is impeded to fully stretch backwards. Furthermore, it is essentially uncoupled from *θ ≈* 2.25rad to *θ ≈ π/*4, and only in the last part is captured by the motor domain. (iv) The swing of MVI lever arm is counterclockwise, as monitored by the observer in Fig. 2b, and occurs on the side of actin. (v) The typical time of rotation from the uncoupled state to the R state for a 7.5nm-lever arm is about 0.2ms. This is consistent with a scheme in which short steps are taken whenever the trailing, free head binds F-actin before the rotation of the uncoupled lever arm is completed, that is before the second step of the PrePS→R transition.

## Conclusions

We studied the power stroke of MVI modeled as the transition from the PrePS state to the R state, focusing on the dynamics associated with the converter, ins2, and IQ domain. Experimental evidence suggests that the converter and ins2 (at least its proximal part) are responsible for the reverse directionality of MVI, and for the large step-size distribution.

Our simulations allow us to draw two major conclusions. (i) The power stroke of MVI occurs in two steps. In the first step, the motor domain undergoes the PrePS→R transition, characterized by the movement of the converter, and then the lever arm rotates to the R-state conformation. (ii) During the rotation, the lever arm is largely uncoupled from the motor domain, which we established using a combination of theory and simulations.

Furthermore, we found that the swing occurs mostly on the side of F-actin, and it is counterclockwise from the perspective of the observer drawn in Fig. 2b-c (see Movie2 and Movie4).

The predictions made are amenable to experimental test.

## Materials and Methods

### Preparation of the PrePS and R states

We prepared the PrePS and R states by combining multiple Protein Data Bank (PDB) structures. The sequence of MVI and the two structures in Fig. 2 show a MVI made of the motor domain, the lever arm up to residue K834 of the IQ domain, and two bound calmodulins (CaMs).

The PrePS and R structures up to residue H786 (located in the middle of ins2, see Fig. 2a) were taken from the PDB structure 4ANJ [41] and 2BKI [34], respectively. Since the remaining of PDB structures 4ANJ and 2BKI were either partially substituted by GFP (4ANJ) or only partially solved (2BKI), we completed our model using the ins2, IQ domain, and related CaMs of the PDB structure 3GN4 [24]. To generate a continuous and seamless model, the residues 776-816 (large part of ins2 and the beginning of the IQ domain, see Fig. 2) of 4ANJ and 2BKI were superposed with the same residues in 3GN4, resulting in a RMSD of 0.38Å and 0.47Å, respectively. The resulting PrePS (R) structure in Fig. 2b (Fig. 2c) was used in our simulations.

Experiments [8, 56] suggest that the lever arm of MVI undergoes the following sequence of transitions: (i) From the ADP and phosphate-bound PrePS state to a conformation in which the lever arm has not moved, but the motor domain is now primed for phosphate release (the PiR state) [56]. (ii) Following the release of phosphate, the MVI head reaches an ADP-bound state. During this transition, most of the swing occurs, but the lever arm conformation does not correspond to the one in the R state [8] (see [20] for a Cryo-EM image). (iii) Following ADP release, the motor finally reaches the R state.

We set PrePS as the initial state, and R as the final state, ignoring the intermediates which are either poorly structurally characterized (ADP-bound), or are close to our initial (PiR) or final (ADP-bound) states.

### Coarse-Grained Model

We created a coarse-grained (CG) representation of MVI, in which each amino acid is represented as a single bead centered at the C_*α*_ position. We used the Self-Organized Polymer (SOP) version of the CG model [57, 58], which is described in the Supplementary Information (SI). Of relevance here is that the nature of the interaction between two beads depends on their distance in the native state. If their distance in the native state is below *R*_C_ = 8Å, and if they are at least 3 residues away from each other in the sequence, the two beads interact through an attractive potential of the Lennard-Jones type, with minimum at their native distance and well depth of *ɛ*_NAT_ = 2kcal*/*mol. The connectivity of the chain is ensured by a finitely-extensible non-linear elastic potential between consecutive beads in the sequence. The interaction between all other pairs of beads is purely repulsive, with strength of the repulsion at the diameter of the beads *σ* = 3.8Å equal to *õ*_NNAT_ = 1kcal*/*mol.

### PrePS→R Transition

To simulate the PrePS→R transition, we adopted the method in [58] with minor variations, which were consistent with the results of fully-atomistic molecular dynamics simulations of MVI carried out using the special purpose Anton computer. This method (see SI for details) uses an energy function that simultaneously incorporates the native interactions in PrePS and in R states, and it mimics the effect of phosphate release by making the interactions associated with the PrePS state weaker, thus facilitating the PrePS→R transition. Recently, a different CG model of MVI illustrated that the release of the phosphate from the nucleotide binding site switches the peak of the population of MVI conformation towards the post-stroke state [16].

We generated 100 trajectories for long enough times to ensure that MVI has completed the PrePS→R transition. For the purpose of data analysis, the overall rotation of MVI was removed by aligning the motor domain before the converter (residues 1-703, see Fig. 2) to the motor domain of the PrePS structure using the Kabsch algorithm [59, 60]. The rotation of the lever arm was monitored after carrying out this alignment. The converter domain and the lever arm, which are subject to the largest change, were not considered in the alignment.

We also generated 100 trajectories with backward loads of 2pN, 4pN, and 6pN. In these cases, to avoid the rigid body movement of MVI due to the external force, we mimic the effect of F-actin by restraining the position of some residues of the motor domain (see SI for details), thus we did not perform any alignment before monitoring the rotation of the lever arm.

Finally, we carried out simulations of MVI in R state and pulled it backwards, so to induce the reverse rotation towards the PrePS state. The pulling forces were 1pN, 10pN, and 20pN. Also in this case, some residues of the motor domain were restrained to avoid the rigid body rotation of MVI under load.

The data used in this paper is available upon request.

## Acknowledgements

MLM is indebted to Yonathan Goldtzvik for numerous conversations about molecular motors and the implementation of the computational models adopted in this study. We are grateful to Matthew Caporizzo, Yale Goldman, Naoto Hori, Xin Li, Sumit Sinha, and Huong Vu for carefully reading the manuscript and providing helpful comments. Anton computer time was provided by the Pittsburgh Supercomputing Center (PSC) through Grant R01GM116961 from the National Institutes of Health. The Anton machine at PSC was generously made available by D.E. Shaw Research. MLM is grateful for the help provided by the PSC team in setting up the simulations on Anton. This work was supported by a grant from the National Science Foundation through grant numbers CHE 16-36424 and CHE 16-32756.

